# Event-related dynamics of brain oscillations during perception of verbal information in patients with paranoid schizophrenia

**DOI:** 10.1101/853705

**Authors:** Zhanna V. Garakh, Ekaterina V. Larionova

## Abstract

We studied the evoked changes in brain rhythmic activity during reading semantic (words) and meaningless verbal information (pseudowords) in patients with paranoid schizophrenia (n= 40) and in control group of healthy subjects (n= 64). Patients with paranoid schizophrenia showed the decrease in the event-related synchronization (ERS) of alpha and theta rhythms compared to controls when reading semantic verbal information. Reduced event-related synchronization of the alpha rhythm in time window 105 – 145 ms may be associated with the severity of hallucinations (P3 scale) by PANSS. In contrast to the control group, the patients had no differences in the gamma range when reading words and pseudowords. This fact may indicate that patients with paranoid schizophrenia assign significance to those stimuli that are not normally considered as significant (pseudowords).

## Introduction

Language disorders are a characteristic feature of schizophrenia. Behavioral and neurobiological experiments demonstrated violations of verbal information processing from the level of single words to text fragments, these violations were found before or during the first psychotic episode and in people at risk of schizophrenia (Goldberg et al., 1998; Kuperberg et al., 2008; Tan et al., 2016; Pawełczyk et al., 2018). It was shown that linguistic abnormalities in schizophrenia may be associated with deficits in automatic and controlled semantic search, attention and working memory (Li et al., 2018; Landre, Taylor, 1995).

Neurophysiological studies show that abnormalities in oscillatory activity may play a crucial role in the pathophysiology of schizophrenia, because neural oscillations are fundamental mechanism of establishing precise time relationships between the reactions of neurons, which, in turn, are important for memory, perception and consciousness (Uhlhaas, Singer, 2015).

We suggest that abnormalities in the synchronized oscillatory activity of neurons during semantic categorization of visually presented verbal information may be due to the mechanisms of schizophrenia. Knowledge about spectral, spatial and temporal features of brain oscillations can help to understand the physiological basis of language and social disorders in schizophrenia. Much attention is paid to the deficiency of early stages of sensory processing in patients with schizophrenia (Martinez et al., 2008). Impaired reading processes in schizophrenia may also be associated with dysfunction of the early stages of information processing (Revheim et al., 2008).

The purpose of this study was to analyze the frequency-time (200 ms after the stimulus) and topographical features of the induced rhythmic activity of the brain during reading semantic and meaningless verbal information in mentally healthy subjects and in patients with paranoid schizophrenia.

## Methods and materials

This study consisted of two groups of participants: 64 healthy subjects, age 27.7±8.1 years (control group) and 40 patients with diagnosis of paranoid schizophrenia, age 28.7±9.7 years (F20.0, experimental group). Diagnosis was based on ICD-10 and was confirmed in a 1-year follow-up. Patients took part in the study 3-20 days after getting to the Department of the first episode of the disease of Moscow Research Institute of Psychiatry. All patients were on anti-psychotic therapy with an average duration of 1 week. Symptom severity were measured using the Positive and Negative Syndrome Scale (PANSS) (Kay et al., 1987).

All participants were right-handed. The overall exclusion criteria were: brain injury, comorbid neurological disorders, somatic illnesses compromising the central nervous system, or an active diagnosis of substance abuse. There were no differences between the groups in gender (p=0.98, t-test) and age (p=0.60, t-test). All participants signed the informed consent.

The subjects were instructed to read the words and pseudowords presented on the screen and not to say them out loud. The stimuli (80 words and 80 pseudowords) were presented in a pseudorandom order on the screen of a 14-inch monitor. All words were nouns and consisted of 5-6 letters. Each stimulus was presented for 100 ms. The interstimulus interval varied from 1500 to 4000 ms. We used the paradigm of simple categorization of single words (word – pseudoword), without the instructions.

Electroencephalogram was recorded from 19 electrodes: Fp1, Fp2, F3, F4, F7, F8, C3, C4, T3, T4, T5, T6, P3, P4, O1, O2 and midline sites (Fz, Cz, Pz) according to the "International 10–20 System". Reference electrodes were placed on the linked earlobes. Low-frequency filters were set at 70 Hz, with a time constant of 0.3 s and the sampling rate of 200 Hz. All electrode impedances were maintained at or below 10 kΩ, with most EEG sites near 5 kΩ. ERP were recorded by a 24-channel amplifier (Medicine-Biology-Neurophysiology, Russia) and 2 synchronized computers to deliver stimuli and to record the ERPs.

For each stimulus type were analyzed changes in event-related synchronization relative to prestimulus interval (300 ms): in the ranges of alpha (8 – 13 Hz) in time window 105 – 145 ms after stimulus, in the ranges of theta (4 – 7.9 Hz) in time window 130 – 210 ms and in the ranges of gamma (30 – 40 Hz) in time window 180 – 230 ms. EEG is filtered digitally within the desired frequency band. Instantaneous values of filtered EEG are squared (to obtain power) and averaged across the selected epochs. Average powers are normalized by the mean power over the base interval (300 ms before the stimulus onset) and transformed to decibels (P_rel_(dB)=10*lg(P_inst_/P_base_), where P_inst_ – instantaneous power, P_base_ – mean baseline power).

We investigated differences in reading between groups. To identify the categorical differences between the stimuli, we compared ERS values in each group of subjects separately. Statistical analysis was done using analysis of variance (ANOVA), the Fisher LSD criterion was applied for post-hoc comparison. Greenhouse–Geisser corrections were applied to correct for violations of sphericity and homogeneity. Correlations between ERS values and PANSS scores were obtained for patients using Pearson correlation coefficients.

## Results and discussion

During reading words and pseudowords all subjects had a same topography of event-related dynamics of brain oscillations in different frequency bands (Fig. 1).

**Figure 1.**
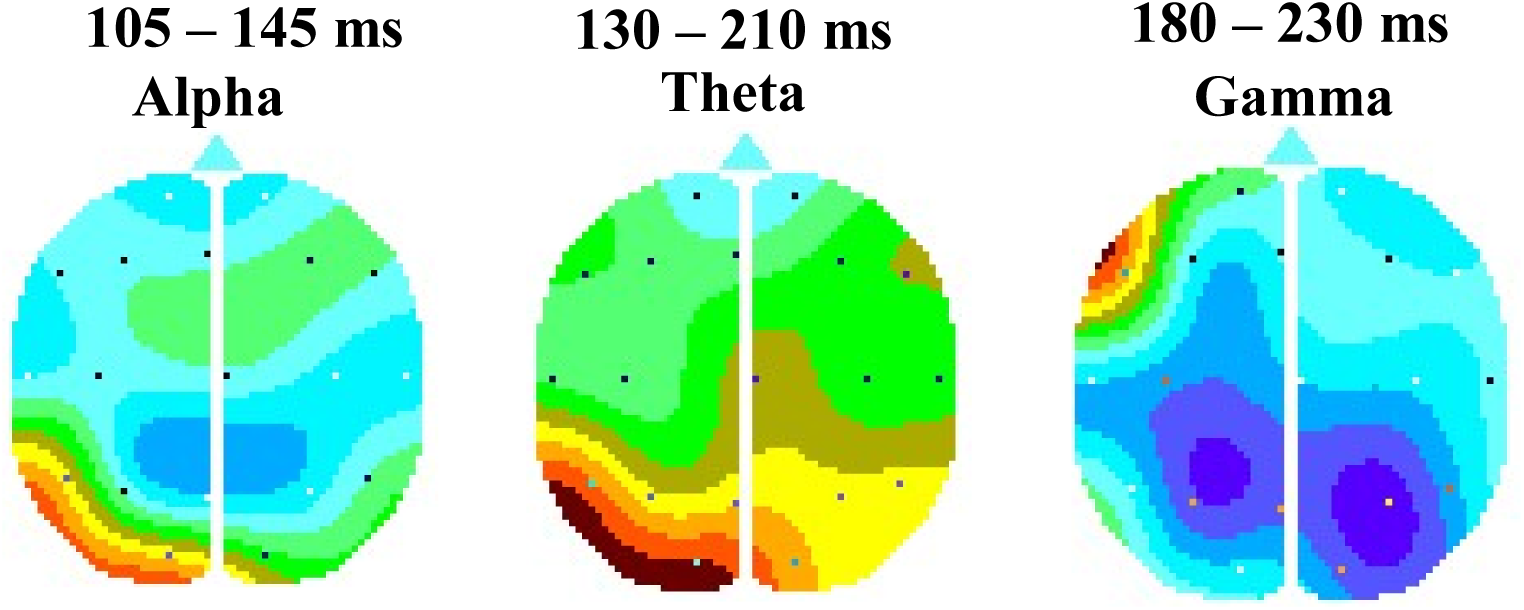
ERS dynamics in reading words in healthy subjects. The scale is given in db. Min – dark blue (ERD), max – red (ERS).

At the first stage (around 125 ms after the stimulus), the synchronization of alpha rhythm in the left occipital, parietal and posterior temporal areas (projection of the "visual word form area") was obtained. The event-related synchronization of the alpha rhythm during reading words was greater in healthy subjects than in patients with schizophrenia (the main “Group” effect, F(1, 102) = 4.62, p=0.03). Post-hoc test showed significant intergroup differences for sites O1 (p<0.01) and T5 (p<0.05). In the earlier studies there was a significant decrease in the synchronization of resting-state alpha in patients with schizophrenia compared to the control group and this decrease was correlated with visual memory scores (Ikezawa et al., 2011). The authors suggest that this decrease in the synchronization of alpha may indicate dysfunction of the left temporal region and impaired visual information processing in schizophrenia, and may also represent a neurophysiological marker of this disease. The detection of alpha-synchronization decrease in the posterior temporal region, especially in the left hemisphere, confirms previous studies indicating a close relationship between left-sided functional and structural abnormalities and the pathophysiology of schizophrenia (Hajek et al., 1997). Dysfunction of the left posterior middle temporal cortex, in particular, may be associated with clinical features in schizophrenia (Job et al., 2002; Whalley et al., 2009). In our study, reduced event-related synchronization of the alpha rhythm in time window 105 – 145 ms in patients correlated with the severity of hallucinations (positive syndrome P3 by PANSS) at a trend level (p=0.06).

In time window 130 – 210 ms during reading words and pseudowords the event-related synchronization of theta rhythm in the left occipital and posterior temporal areas was obtained. The event-related synchronization in this time window during reading words and pseudowords was greater in healthy subjects than in patients with schizophrenia in the occipital O1 (p<0.01), and parietal-temporal areas T5 (p<0.05). Decrease in the synchronization of theta rhythm in patients with schizophrenia was shown during the problem of recognition of facial expression (Csukly et al., 2014, Marosi et al., 2019). Categorical differences between word and pseudoword in ERS values were found neither in healthy subjects nor in patients. The subjects of the two groups showed no differences in the event-related synchronization of alpha and theta rhythms during reading semantic and meaningless verbal information.

In time window 180 – 230 ms during reading words and pseudowords the event-related synchronization of gamma rhythm in anterior temporal electrode (F7, Broca’s area) of the left hemisphere was obtained. There were no intergroup differences in event-related synchronization. The healthy subjects had differences in the gamma range when reading words and pseudowords: ERS of gamma was greater when reading words than pseudowords (p<0.0001). This time interval can be associated with a lexical access (Almeida et al., 2013) and stimulus categorization process (Pernet et al., 2003). In particular, a power increase in the gamma frequency band was demonstrated in the perception of words related to the meaning of the proposed sentence (Rommers et al., 2013). In contrast to the control group, the patients with paranoid schizophrenia had no differences in the gamma range when reading words and pseudowords.

## Conclusion

In this study we found abnormalities in the synchronized oscillatory activity of neurons during reading in patients with paranoid schizophrenia. At the early stages of sensory analysis of visually presented verbal information, patients with paranoid schizophrenia showed the decrease in the event-related synchronization (ERS) compared to controls only when reading semantic information (words). At the stage of lexical decision (200 ms after stimulus) in contrast to the control group, the patients had no differences in the gamma range when reading words and pseudowords. This fact may indicate that patients with paranoid schizophrenia assign significance to those stimuli that are not normally considered as significant (pseudowords).

## Acknowledgements

This study was partially supported by the RFBR (grant № 18-013-00733) and funds within the state assignment of Ministry of Education and Science of the Russian Federation for 2019-2021 (No. AAAA-A17-117092040004-0).

## Disclosure

No potential conflict of interest.

